# Hepatocytic Prominin-1 protects against liver fibrosis by stabilizing the SMAD7 protein

**DOI:** 10.1101/846493

**Authors:** Hyun Lee, Dong-Min Yu, Min Jee Um, Seo Yeon Yoon, Ki-Tae Kim, Young Jae Kwon, Sungsoo Lee, Myeong-Seok Bahn, Seung-Hoi Koo, Ki Hoon Jung, Jae-Seon Lee, Young-Gyu Ko

**Author notes:** These authors contributed equally (Hyun Lee, Dong-Min Yu). To whom correspondence should be addressed Young-Gyu Ko, Ph. D. Division of Life Sciences, Korea University 145, Anam-ro, Seongbuk-gu, Seoul, 02841, Korea. **Author contributions** H.L., D.-M.Y., M.J.U., S.Y.Y., K.-T.K., S.S.L. Y.J.K., and M.-S.B. performed the experiments; K.H.J. collected the patient liver specimens; S.H.K. created liver-specific knockout; J.-S.L. and Y.-G.K. designed the experiments and analyzed the data; and H.L., D.-M.Y. and Y.-G.K. wrote the manuscript.

## Abstract

**Background and Aims:** Prominin-1 (PROM1) is known to be upregulated in hepatocytic progenitor cells (HPCs) and cholangiocytes of fibrotic livers. To understand the function of upregulated PROM1 during liver fibrosis, we analyzed liver fibrosis from global and liver-specific Prom1 knockout mice and investigated the molecular mechanism of how Prom1 protects the liver against liver fibrosis.

**Methods:** We analyzed PROM1 expression from human liver with mild and severe liver fibrosis (n=4-9). Liver fibrosis was induced by carbon tetrachloride (CCl_4_) treatment and bile duct ligation (BDL) from wild type and global and liver-specific knockout mice (n=3-13). The severity of liver fibrosis was determined by qRT-PCR, immunostaining and immunoblotting for fibrotic markers such as αSMA, collagen. TGFβ signaling was also analyzed from fibrotic liver and primary hepatocytes of wild type and global and liver-specific knockout mice (n=3-5). Molecular interaction between PROM1 and SMAD7 was determined by endogenous and exogenous co-immunoprecipitation.

**Results:** PROM1 was found in the plasma membranes of both healthy and fibrotic hepatocytes and cholangiocytes. Global Prom1 knockout aggravated BDL- and CCl_4_-induced liver fibrosis. *Prom1*^-/-^ hepatocytes showed increased TGFβ signaling due to reduced SMAD7 protein expression compared to that in wild-type hepatocytes. PROM1 prevented SMURF2-induced SMAD7 ubiquitination and degradation by interfering with the molecular association of SMAD7 with SMURF2. We also demonstrated that liver-specific Prom 1 knockout aggravated BDL-induced liver fibrosis due to reduced levels of SMAD7.

**Conclusion:** Hepatocytic PROM1 stabilizes SMAD7, preventing TGFβ signaling. Thus, PROM1 is necessary for the negative regulation of TGFβ signaling during liver fibrosis.

## Introduction

Liver fibrosis results from the accumulation of collagen scar tissue following chronic liver injuries caused by nonalcoholic steatohepatitis (NASH), alcohol, hepatitis viral infection and autoimmune hepatitis ^1, 2^. Chronic liver fibrosis progresses into hepatocellular carcinoma and cirrhosis with characteristics of ascites, jaundice (yellow skin), hepatic hypertension and hepatic encephalopathy if the cause is not eliminated ^3^. Liver fibrosis is initiated by damaged hepatocytes and infiltrated macrophages, which secrete profibrogenic cytokines such as transforming growth factor-β (TGFβ), platelet-derived growth factor (PDGF), connective tissue growth factor (CTGF), vascular endothelial growth factor (VEGF), sonic hedgehog (SHH) and WNT4. Hepatic stellate cells (HSCs) residing between hepatocytes and endothelial cells are proliferated and activated by these profibrogenic cytokines to produce extracellular matrix (ECM) components such as collagen and fibronectin ^4–6^.

TGFβ binds to its type II receptor (TβRII), recruiting and phosphorylating its type I receptor (TβRI) in HSCs. Activated TβRI induces the phosphorylation of SMAD2 and SMAD3, and the activated SMAD2/3 then binds to SMAD4. The SMAD2/3-SMAD4 complex enters the nucleus and acts as a transcription factor to upregulate the genes encoding type I collagens (*COL1A1* and *COL1A2*), tissue inhibitors of matrix metalloproteases (*TIMPs*) and α-smooth muscle actin (*ACTA2*) ^7, 8^. The activated SMAD2/3-SMAD4 complex also upregulates SMAD7, a negative feedback inhibitor of TGFβ signaling. SMAD7 binds to TβRI and induces the ubiquitination and degradation of TβRI by recruiting SMAD ubiquitination regulatory factors 1 and 2 (SMURF1 and 2), thereby preventing TGFβ-induced phosphorylation and activation of SMAD2/3 ^9–11^. SMAD7 also prevents the phosphorylation of SMAD3 or formation of the SMAD2/3-SMAD4 complex without TβRI ubiquitination ^12, 13^.

TGFβ plays important roles in hepatocytes during liver fibrosis ^7, 8^. For example, hepatocyte-specific *Tgfbr2* deletion or *Smad7* overexpression prevents liver fibrosis development via NASH, bile duct ligation (BDL) and carbon tetrachloride (CCl_4_) ^14–16^. Transdifferentiation of primary-cultured hepatocytes into myofibrils is also prevented by adenoviral overexpression of *Smad7* ^14^.

Prominin-1 (PROM1), also called CD133, is a pentaspan transmembrane glycoprotein expressed in hematopoietic, epithelial and intestinal progenitor cells ^17, 18^. PROM1 is expressed in rat HSCs and has been used as a cell-surface marker to isolate hepatic progenitor cells (HPCs) ^19^. High expression levels of PROM1 are also observed in HPCs and cholangiocytes after liver injuries, such as those resulting from rhesus rotavirus (RRV)-induced biliary atresia (BA) and BDL ^20, 21^. Because Prom1 deficiency was shown to protect against RRV-induced liver fibrosis in mice, upregulated Prom1 in HPCs might be necessary to promote biliary fibrosis ^22^.

In addition to HPCs and cholangiocytes, we found PROM1 to be expressed in hepatocyte plasma membranes of both healthy and fibrotic livers. We also demonstrated that PROM1 deficiency aggravated BDL- and CCl_4_-induced liver fibrosis and enhanced TGFβ signaling by reducing SMAD7 protein expression in hepatocytes. Because PROM1 interfered with the molecular association of SMAD7 with SMURF2 and prevented SMURF2-induced SMAD7 ubiquitination and degradation, we concluded that PROM1 is necessary for the prevention of liver fibrosis by negatively regulating TGFβ signaling.

## Materials and methods

### Patient samples

Samples from humans with mild or severe fibrosis were obtained from Dongguk University in Gyeongju, South Korea. The patients had been diagnosed with liver fibrosis or cirrhosis by histological examination at hospitals in South Korea.

### Animal models and experiments

Prom1 knockout mice (The Jackson Laboratory, Bar Harbor, ME, USA), in which the ATG start codon of the CD133 gene was replaced by a CreERT2 fusion protein and an IRES-b-galactosidase (lacZ) gene, were utilized. Prom1 KO mice were backcrossed onto a C57BL/6N background for at least five generations.

To generate liver-specific knockout mice of Prom1 (*f/f : Alb-Cre*), Prom1^loxP/loxP^ (*f/f*) mice were created by ToolGen (Seoul, Korea) and crossed with Alb-Cre (*Alb-Cre*) mice containing a *Cre* recombinase driven by albumin promoter (The Jackson Laboratory, Bar Harbor, ME, USA).

Mice, housed in plastic cages under a 12:12 light-dark photoperiod with free access to water and food, were bred, maintained, and cared for in a manner consistent with criteria outlined in the Principles of Laboratory Animal Care (NIH publication no. 85-23, revised 1985), and protocols were approved by the Institutional Animal Care and Use Committee of Korea University. Notably, CD133 KO mice are viable and live to adulthood without exhibiting phenotypic differences from their WT littermates.

### Mouse model of liver fibrosis

To generate different mouse fibrosis models, CCl_4_ and BDL treatment were performed. Toxicant-induced models were generated by subcutaneous injections with either CCl_4_ (1 ml/kg body weight) in corn oil (1:3 dilution), as a control, in six-week old male WT and KO C57BL/6 mice every third day for 6 weeks. The mice were killed 2 days after the final CCl_4_ injection.

For the BDL model, the mice were randomly divided into two groups. Animals that underwent BDL and sham operated animals were used as healthy controls. Briefly, mice were anesthetized using isoflurane. The extra-hepatic bile duct was isolated and doubly ligated. The peritoneal cavity was filled with saline and the incisions were closed. Mouse body weight were measured daily. Seven days after the surgery, BDL and sham control animals were sacrificed.

### Serum biochemistry

Serum ALT, AST and total bilirubin levels were analyzed using spectra-iMAX (Molecular Devices) and commercial kits from Biovision (San Francisco, USA). Serum levels were analyzed using commercial kits according to the manufacturer instructions.

### Immunohistochemistry

Freshly frozen tissues post-fixed in 4% paraformaldehyde were subjected to immunohistochemistry analysis. Tissue sections were immunostained with antibodies against prominin-1 (eBioscience or Milteny Biotec), CK19, αSMA, Smad7, p-Smad2/3, Smad 2/3 and Vimentin. Briefly, the sections were pretreated with 2.5% horse serum for 20 min to block nonspecific antibody binding and then incubated with the antibodies of interest for overnight at 4°C. The slides were then treated with a FITC- or rhodamine-conjugated secondary antibody. After mounting with Permount solution, the samples were imaged on an LSM 800 META confocal microscope (Carl Zeiss, Thornwood, NY).

### Preparation of mouse primary hepatocytes

Primary hepatocytes were isolated from 8-week-old C57BL/6 male mice as previously described. Briefly, mice were anesthetized with avertin (intraperitoneal injection of 250 mg/kg body weight), and livers were perfused with a pre-perfusion buffer (140 mM NaCl, 6 mM KCl, 10 mM HEPES, and 0.08 mg/mL EGTA, pH 7.4) at a rate of 7 mL/min for 5 min, followed by a continuous perfusion with a collagenase-containing buffer (66.7 mM NaCl, 6.7 mM KCl, 5 mM HEPES, 0.48 mM CaCl_2_, and 3 g/mL collagenase type IV, pH 7.4) for 8 min. Viable hepatocytes were harvested and purified with a Percoll cushion. Then, hepatocytes were resuspended in complete growth medium (199 medium containing 10% FBS, 23 mM HEPES and 10 nM dexamethasone) and seeded on collagen-coated plates at a density of 300,000 cells/ml. After a 4-h attachment period, the medium was replaced with complete growth medium before being used in any experiments and were changed daily.

### Immunofluorescence staining

Livers and cells were fixed in 4% paraformaldehyde, and staining was performed as described previously. Signals were detected by secondary antibodies conjugated to either with anti-mouse, anti-rat rhodamine, or anti-rabbit antibodies. Fluorescence images were acquired on an LSM800 confocal microscope controlled by ZEN software (Carl Zeiss, Thornwood, NY). Bright-field images were acquired using a transmitted light microscope (Leica microsystems AG, Heerbrugg, Switzerland) and the Leica application system.

### Western blotting analysis

Immunblot analyses were performed as previously described. Briefly, cells were harvested with lysis buffer (25 mM HEPES, 150 mM NaCl, 1% NP-40, 10 mM MgCl2, and 1 mM EDTA, pH 7.5) on ice for 20 min. The lysates were centrifuged at 10,000 g for 10 min to obtain supernatants. Proteins were separated by SDS–polyacrylamide gel electrophoresis, and the immobilized proteins were immunoblotted with the antibodies of interest. The antigens were visualized using an ECL substrate kit (Thermo Scientific, USA).

For immunoprecipitation, cells were lysed in buffer containing 20 mM Tris-HCl (pH 7.4), 137 mM NaCl, 1 mM MgCl_2_, 1 mM CaCl_2_ and a protease inhibitor cocktail (Sigma-Aldrich, St Louis, MO, USA). Whole-cell lysates (500 µg of protein) were incubated with specific antibodies overnight and then with 60 μl of slurry of Protein A- or Protein G-agarose beads (Roche, Mannheim, Germany) for 3 h. Immunoprecipitants were analyzed by immunoblot.

### Quantitative real-time PCR

RNA (2 μg) was reverse transcribed into cDNA using random hexamer primers, oligo dT and reverse transcription master premix (ELPIS Biotech, Daejeon, Korea). Quantitative real-time PCR analyses were performed using the cDNAs from the reverse transcription reactions and gene-specific oligonucleotides in the presence of TOPreal qPCR 2X premix (Enzynomics, Daejeon, Korea). The following PCR conditions were used: an initial denaturation step at 95°C for 10 min, followed by 45 cycles of denaturation at 95°C for 10 s, annealing at 58°C for 15 s and elongation at 72°C for 20 s. The melting curve for each PCR product was assessed for quality control. Supplementary Table 2 shows the sequences of the primers used for qPCR.

### Statistical analysis

Statistical values are presents as the mean ± S.E.M. A two-t-test was used to compare groups. *P<0.05, ** P <0.01, *** P <0.001.

## Results

### PROM1 is expressed in hepatocytes and CK19-expressing cells of patients with liver fibrosis

The *PROM1* mRNA levels in 40 cirrhotic and 6 healthy livers from a human GEO database (GEO accession code GSE25097) were analyzed. Compared to those in the healthy livers, the mRNA levels of both *PROM1* and fibrogenic genes such as those encoding α-smooth muscle actin (*ACTA2*), α1 type 1 collagen (*COL1A1*), and transforming growth factor β1 (*TGFB1*) were upregulated in the fibrotic liver, while mRNA level of *TGFBR1* was not (Fig. 1A). The *PROM1* mRNA levels were increased in proportion to those of *ACTA2, COLA1,* and *TGFB1* in fibrotic livers (Fig. 1B), indicating that the *PROM1* expression level is positively correlated with the progression of liver fibrosis. Next, we determined which cells express PROM1 in livers with mild and severe fibrosis by double immunofluorescence staining for cytokeratin-19 (CK19 for cholangiocytes and HPCs) and α-smooth muscle actin (αSMA for HSCs). PROM1 was mainly found in CK19-expressing cells but not in αSMA-expressing cells (Fig. 1C), indicating that PROM1 is expressed in the cholangiocytes and HPCs of the human fibrotic liver. PROM1 was also clearly observed in hepatocyte plasma membrane (Fig. 1C).

**Figure 1.**
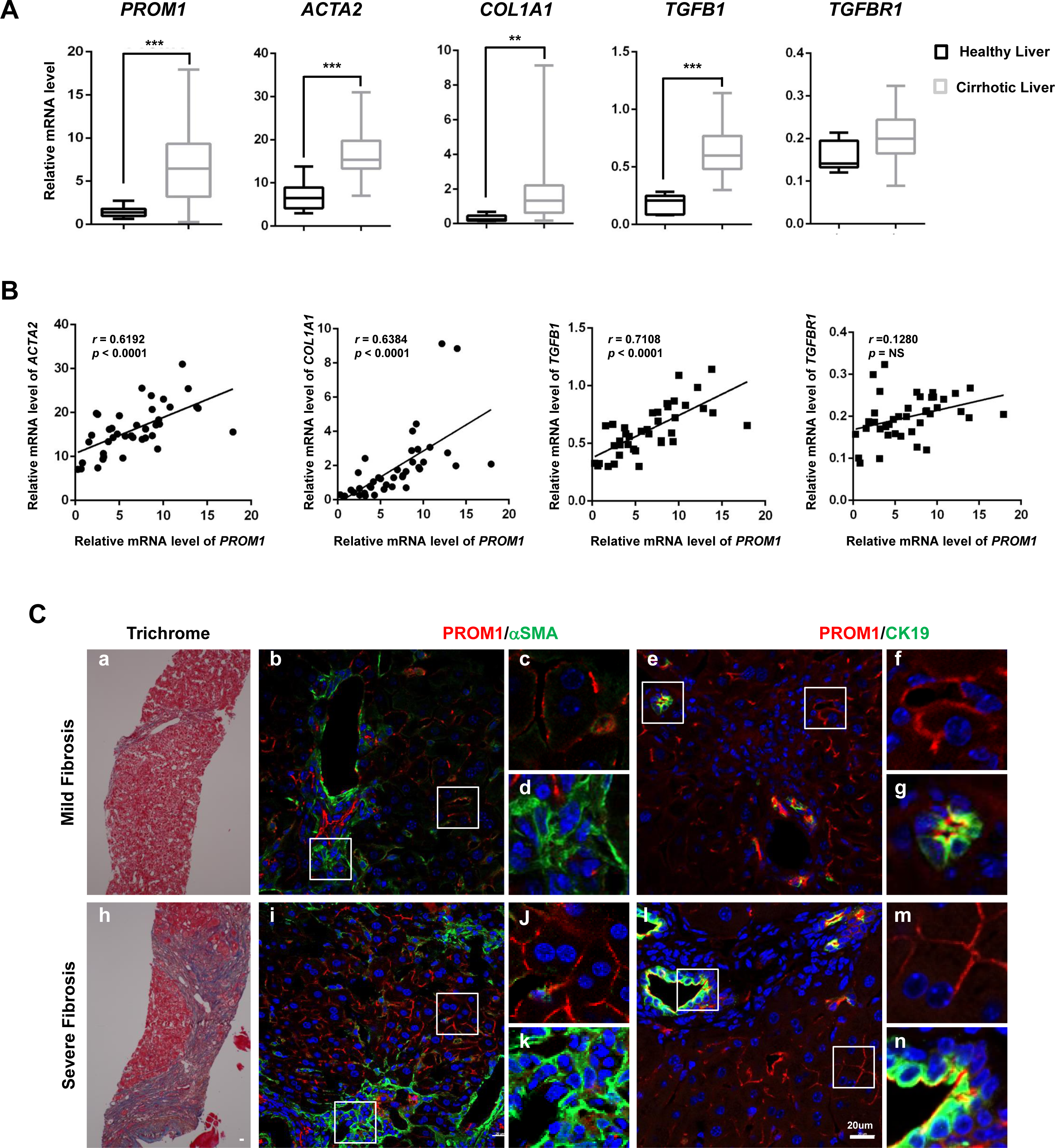
PROM1 is found in both the hepatocytes and CK19-expressing cells of patients with liver fibrosis. (A and B) The relative mRNA levels of *PROM1* and fibrogenic genes (*COL1A1, ACTA2, TGFB,1 and TGFR1*) in 40 cirrhotic and 6 healthy livers were obtained from a human GEO database (GEO accession code GSE25097) and presented using a box-whisker plot (A). Correlations of the relative mRNA levels of *PROM1* and fibrogenic genes were analyzed by Pearson’s *r* test (B). *PROM1*, prominin-1; *COL1A1*, alpha-1 type 1 collagen; *ACTA2*, alpha-actin-2; *TGFB1*, transforming growth factor beta 1; *TGFBR1*, transforming growth factor beta receptor 1. (C) Human liver specimens with mild and severe liver fibrosis were analyzed by Masson’s Trichrome staining (left panel) and immunofluorescence staining for PROM1 (red), αSMA (green) and CK19 (green) (middle and right panels). The specimens were also stained with DAPI (blue). \*\**p* < 0.01, \*\*\**p* < 0.001.

To further confirm the Prom1 upregulation in liver fibrosis, we induced liver fibrosis in mice by BDL and then assessed the liver *Prom1* mRNA levels. In the BDL-treated liver, the *Prom1* mRNA levels were increased in proportion to those of the fibrogenic markers *Acta2, Col1a1, and Tgfb1* but not to the mRNA level of *Tgfbr1* (Fig. 2A). We also determined the tissue distribution of PROM1 by double immunofluorescence staining for CK19 and αSMA in the liver. Prom1 was found in the livers of both sham and BDL mice in all CK19-expressing cells but not in αSMA-expressing cells (Fig. 2B). The livers of BDL mice had substantially more PROM1- and CK19-expressing cells than those of the sham mice (Fig. 2B). These results indicate that Prom1 upregulation in the liver of BDL mice results from the expansion of CK19- and PROM1-expressing HPCs and cholangiocytes. Prom1 was also found in the plasma membrane of hepatocytes, while its expression in the BDL mouse liver was not altered (Fig. 2B).

**Figure 2.**
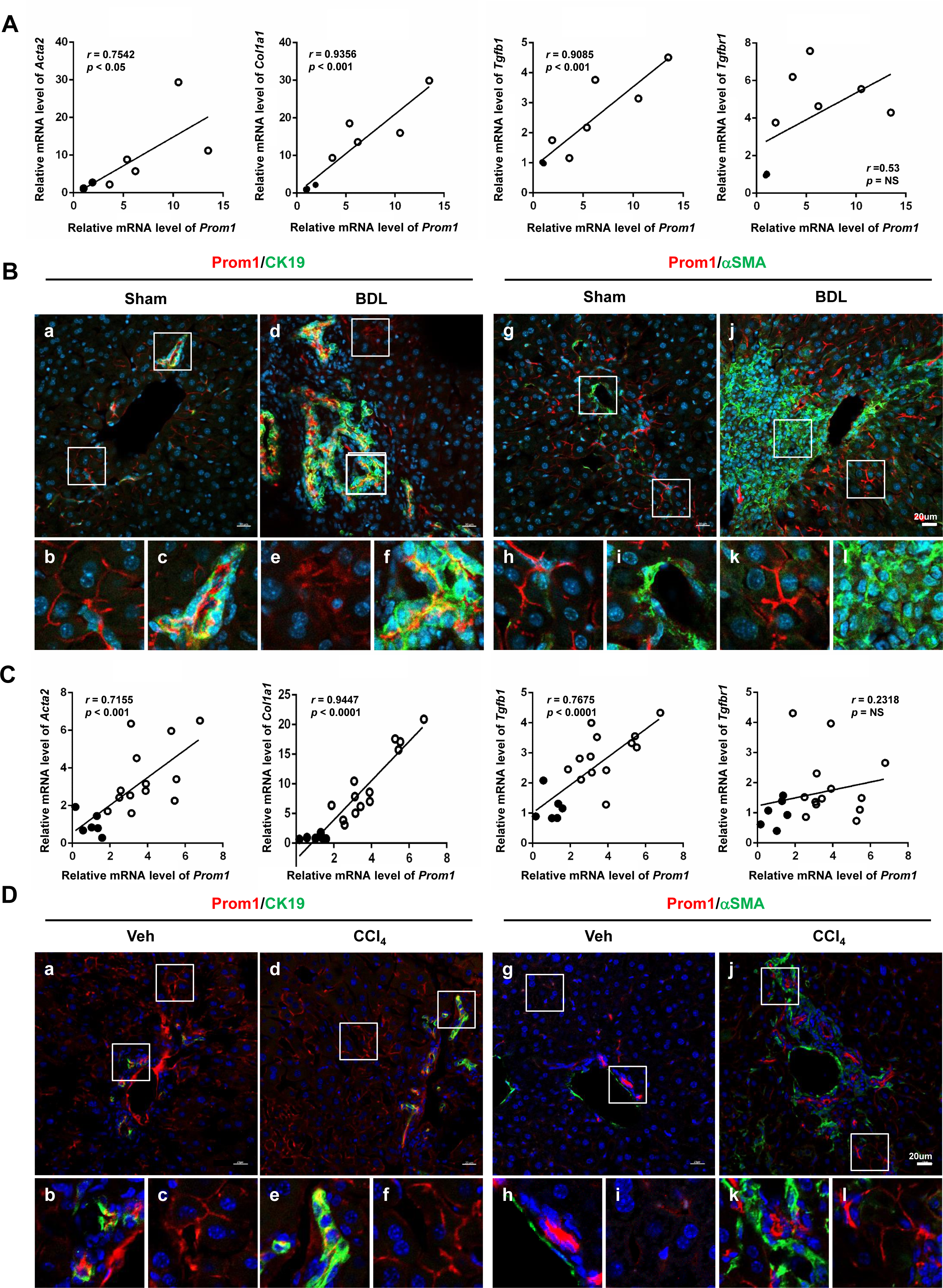
PROM1 is found in the hepatocytes and CK19-expressing cells of mice with liver fibrosis. (A and B) Eight-week-old male mice were subjected to sham (n=3) or BDL (n=6) treatment for one week. The mRNA levels of *Prom1* and fibrogenic genes (*Col1a1, Tgfb,1* and *Tgfbr1*) were determined by qRT-PCR and normalized to those of 18S rRNA. Correlations of the relative mRNA levels between of *PROM1* and fibrogenic genes were analyzed by Pearson’s *r* test (A). The liver specimens were analyzed by immunofluorescence staining for PROM1 (red), αSMA (green), and CK19 (green). Blue represents DAPI-stained nuclei. (C and D) Eight-week-old male mice were administrated an intraperitoneal injection of vehicle (n=6) or CCl_4_ (n=13) twice a week for 6 weeks. The mRNA levels of *Prom1* and fibrogenic genes (*Col1a1, Tgfb1* and *Tgfbr1*) were determined by qRT-PCR and normalized to those of 18S rRNA. Correlation of the relative mRNA levels of *PROM1* and fibrogenic genes were analyzed by Pearson’s *r* test (C). The liver specimens were analyzed by immunofluorescence staining for PROM1 (red), αSMA (green), and CK19 (green) (D). Blue represents DAPI-stained nuclei.

In CCl_4_-treated mice, *Prom1* mRNA levels were also increased in proportion to the mRNA levels of *Acta2, Col1a1, Tgfb1* but not to the mRNA level of *Tgfbr1* (Fig. 2C). Double immunofluorescence analysis showed that PROM1 was expressed in CK19-expressing cells but not in αSMA-expressing cells (Fig. 2D). Prom1 was also clearly expressed in the hepatocyte plasma membrane of both control and CCl_4_-treated mice (Fig. 2D).

### PROM1 deficiency aggravates BDL- and CCl_4_-induced liver fibrosis in mice

To understand the physiological function of PROM1 in the development of liver fibrosis, we induced liver fibrosis in *Prom1*^+/+^ (WT) and *Prom1*^-/-^ (KO) mice by BDL or CCl_4_ treatment. Prom1-expressing cells were found in only the *Prom1*^+/+^ livers and their numbers were increased after BDL and CCl_4_ treatment, indicating that the immunofluorescence signal was specific for PROM1. Hematoxylin and eosin (H&E) staining showed that the *Prom1*^-/-^ liver had more nuclei around the central vein than the *Prom1*^+/+^ liver after BDL or CCl_4_ treatment (Supplementary Fig. 1A). Masson’s trichrome, Sirius Red and αSMA immunostaining showed that BDL- or CCl_4_-induced collagen deposition and myofibroblast differentiation in the liver were increased by Prom1 deficiency (Fig. 3A). However, the number of CK19-expressing cells was significantly increased in *Prom1*^-/-^ mice compared to that in *Prom1*^+/+^ mice in the BDL model but not in CCl_4_-treated mice (Fig. 3A).

**Figure 3.**
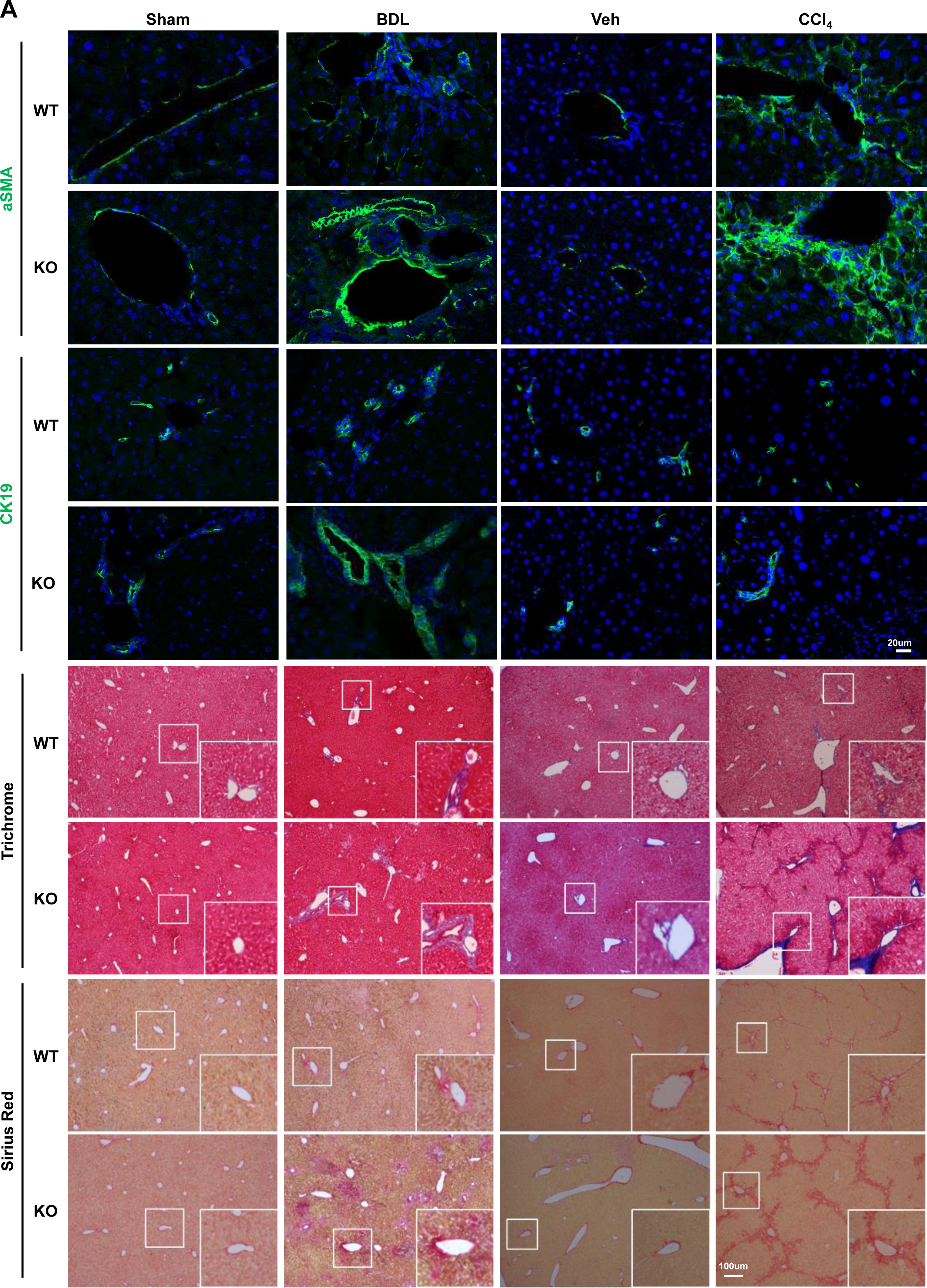
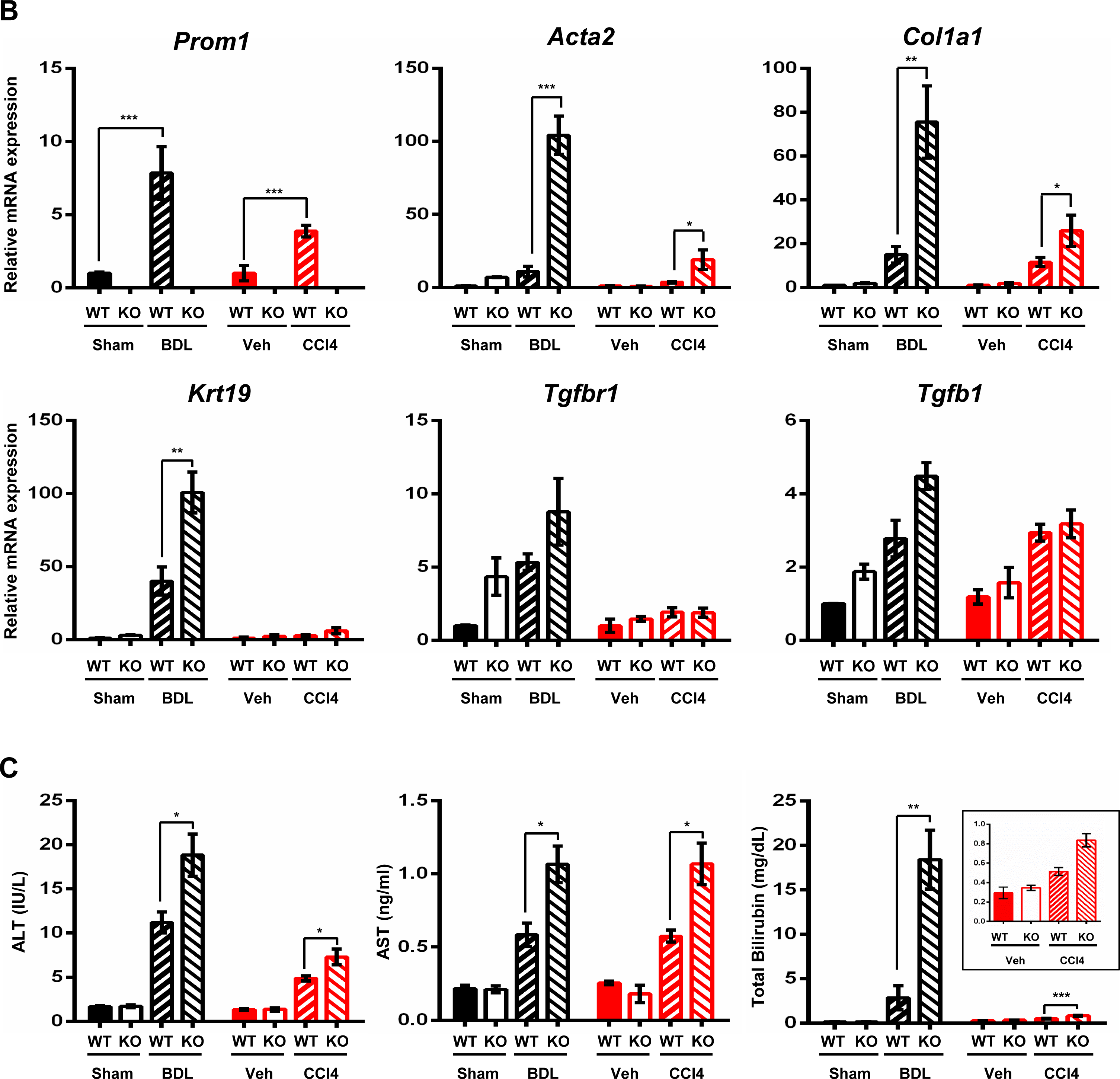
PROM1 deficiency aggravates BDL- and CCl_4_-induced liver fibrosis. For BDL-induced liver fibrosis model, eight-week old male *Prom1*^+/+^ and *Prom1*^-/-^ mice were subjected to sham (n=3) or BDL (n=5-6) treatment for one week. For the CCl_4_-induced liver fibrosis model, eight-week old male *Prom1*^+/+^ and *Prom1*^-/-^ mice were administrated with intraperitoneal injections of vehicle (n=6) or CCl_4_ (n=12-13) twice a week for 6 weeks. Each liver specimen was analyzed by Masson’s Trichrome staining, Sirius Red staining, and αSMA and CK19 immunofluorescence (A). The mRNA levels of *Prom1, Acta2, Col1a1, Krt19, Tgfbr1* and *Tgfbr1* were determined by qRT-PCR and normalized to those of 18S rRNA (B). The serum levels of aspartate transaminase (AST), alanine aminotransferase (ALT), and bilirubin were determined (C). \*\**p* < 0.01, \*\*\**p* < 0.001.

To determine BDL- or CCl_4_-induced liver fibrosis in *Prom1*^+/+^ and *Prom1*^-/-^ mice, we further analyzed the mRNA levels of genes associated with liver fibrosis. In BDL mice, Prom1 deficiency further increased the mRNA levels of TGFβ target genes (*Col1a1, Acta2, Ctgf, Pai1* and *Timp1*), ductular reaction-related genes (*Krt19*), and cytokine genes *(Il1b, Il6, Tnfa,* and *Ccl2*) (Fig. 3B and Supplementary Fig. 1B). However, in CCl_4_-treated mice, although Prom1 deficiency further increased the mRNA levels of TGFβ target genes (*Col1a1*, *Acta2*, and *Timp1*), it did not change the mRNA levels of ductular reaction-related genes (*Krt19*), and cytokine genes *(Il1b, Il6, Tnfa,* and *Ccl2*).

Moreover, the blood levels of aspartate aminotransferase (AST), alanine aminotransferase (ALT) and bilirubin increased by BDL or CCl_4_ treatment were further increased by PROM1 deficiency (Fig. 3C), suggesting that PROM1 deficiency aggravates BDL- or CCl_4_-induced liver fibrosis.

### PROM1 deficiency enhances TGF**β**1 signaling in hepatocytes

Because Prom1 deficiency did not change the mRNA levels of *Tgfb1* and *Tgfbr1* in either BDL or CCl_4_-treated mice, we speculated that Prom1 may regulate TGFβ downstream signal transduction (Fig. 3B). Thus, we first examined TGFβ signaling in the liver after BDL and CCl_4_ treatment. Immunostaining and immunoblotting for phospho-SMAD2/3 (p-SMAD2/3) showed that the BDL- and CCl_4_-induced phosphorylation of SMAD2/3 was increased to a greater extent in the *Prom1*^-/-^ liver, than in the *Prom1*^+/+^ liver (Fig. 4A and B).

**Figure 4.**
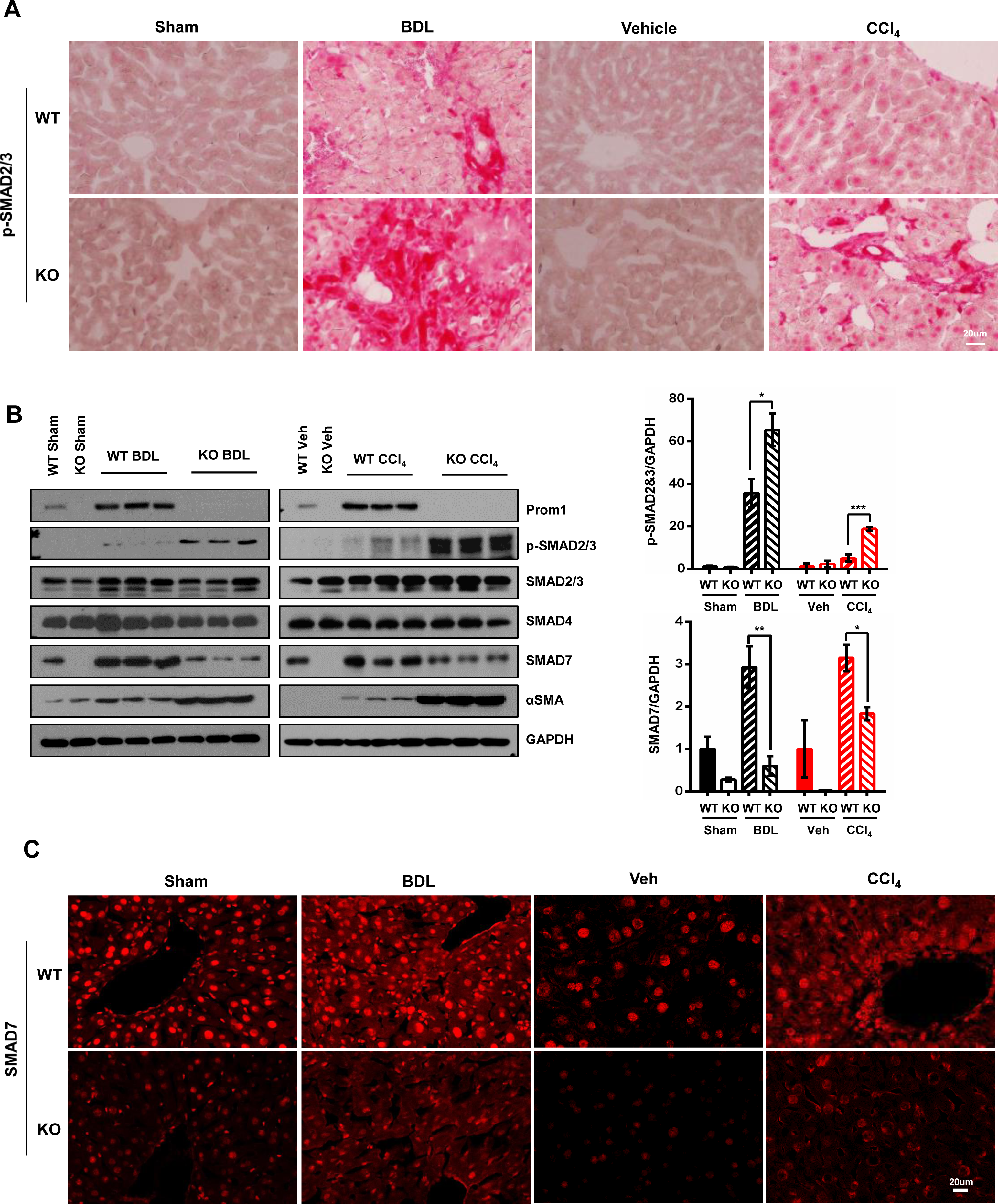
PROM1 deficiency enhances TGFβ signaling by reducing the SMAD7 expression in mice with liver fibrosis. For the BDL-induced liver fibrosis model, eight-week old male *Prom1*^+/+^ and *Prom1*^-/-^ mice were subjected to sham or BDL treatment for one week. For the CCl_4_-induced liver fibrosis model, eight-week old male *Prom1*^+/+^ and *Prom1*^-/-^ mice were administrated with intraperitoneal injections of vehicle or CCl_4_ twice a week for 6 weeks. Each specimen was analyzed by immunofluorescence analysis of phosphorylated SMAD2/3 (p-SMAD2/3) (A) and immunoblotting for PROM1, p-SMAD2/3, total SMAD2/3 (t-SMAD2/3), SMAD4, SMAD7, αSMA, and GAPDH (B, left panel). The expression levels of pSMAD2/3 and SMAD7 were normalized to that of GAPDH and statistically assessed. N=3 for each group. (B, right panel). Each specimen was also subjected to SMAD7 immunofluorescence analysis (C). \**p* < 0.05, \*\**p* < 0.01, \*\*\**p* < 0.001.

Interestingly, Prom1 deficiency prevented the BDL- and CCl_4_-induced increase in SMAD7 protein expression without changing the mRNA level of *Smad7* (Fig. 4B and C and Supplementary Fig. 2A).

Because PROM1 was expressed in hepatocytes and Prom1 deficiency aggravated CCl_4_-induced liver fibrosis in which hepatocytes were mainly damaged, we examined the effect of PROM1 on TGFβ signaling in primary hepatocytes isolated from *Prom1*^+/+^ and *Prom1*^-/-^ mice. *Prom1* deficiency further increased the TGFβ-induced phosphorylation of SMAD2/3 and TGFβ-induced Vimentin (Fig. 5A and B). The mRNA levels of TGFβ target genes (*Snail1*, *Vimentin*, *Zeb1*, and *Zeb2*) were also increased in TGFβ-exposed primary hepatocytes by Prom1 deficiency (Fig. 5C). Interestingly, SMAD7 expression was higher in *Prom1*^+/+^ hepatocytes than in *Prom1*^-/-^ hepatocytes (Fig. 5A and B). Because the mRNA levels of *Smad7* were not decreased in the liver or primary hepatocytes of *Prom1*^-/-^ mice (Supplementary Fig. 2A and 3B), SMAD7 might be posttranslationally stabilized in the presence of PROM1. Next, we examined TGFβ signaling after the adenoviral overexpression of *PROM1* in *Prom1*^-/-^ primary hepatocytes. Immunofluorescence and immunoblot analyses showed that PROM1 restoration prevented the TGFβ-induced phosphorylation of SMAD2/3 expression by increasing SMAD7 protein expression (Fig. 5D and E).

**Figure 5.**
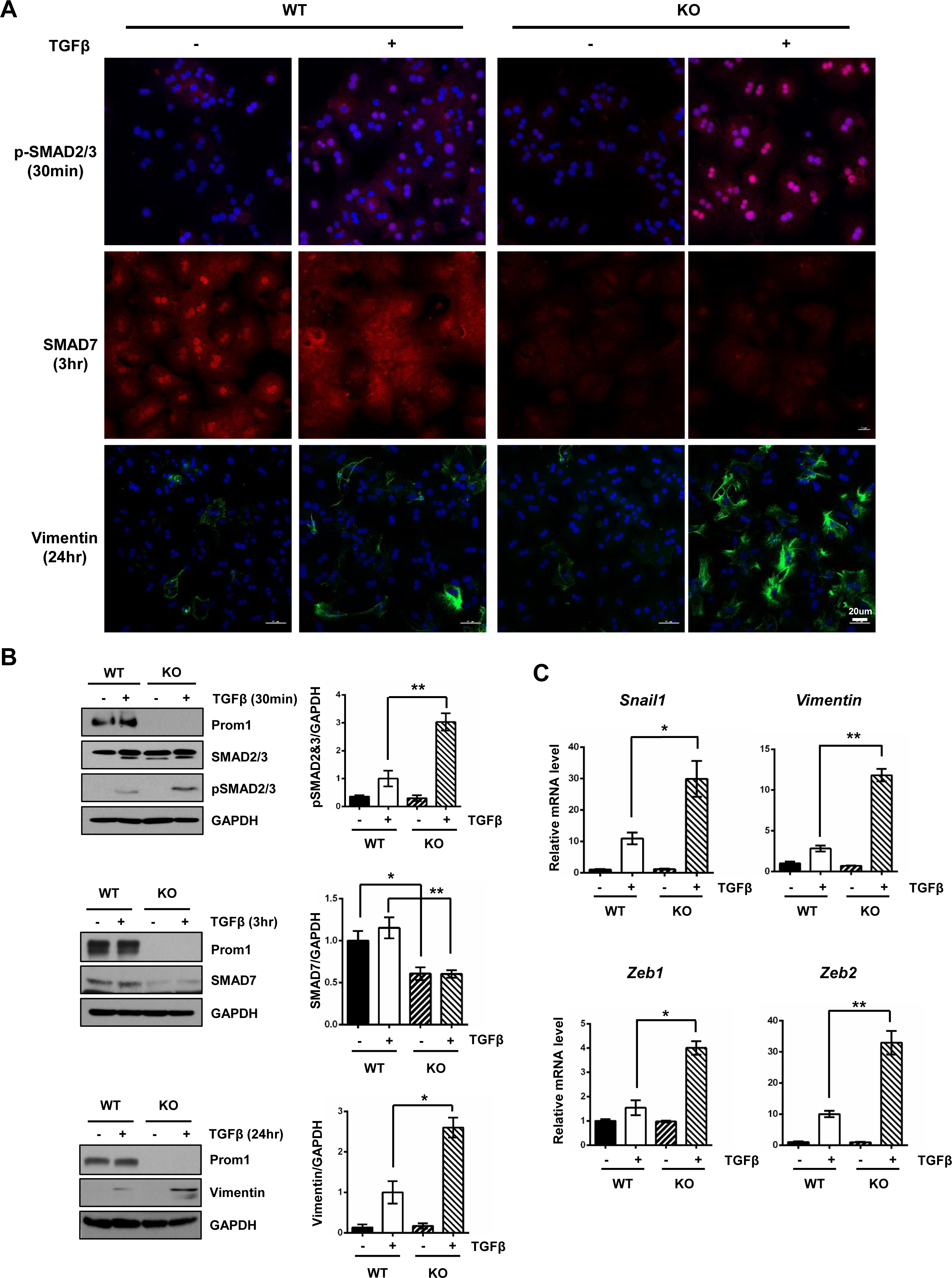
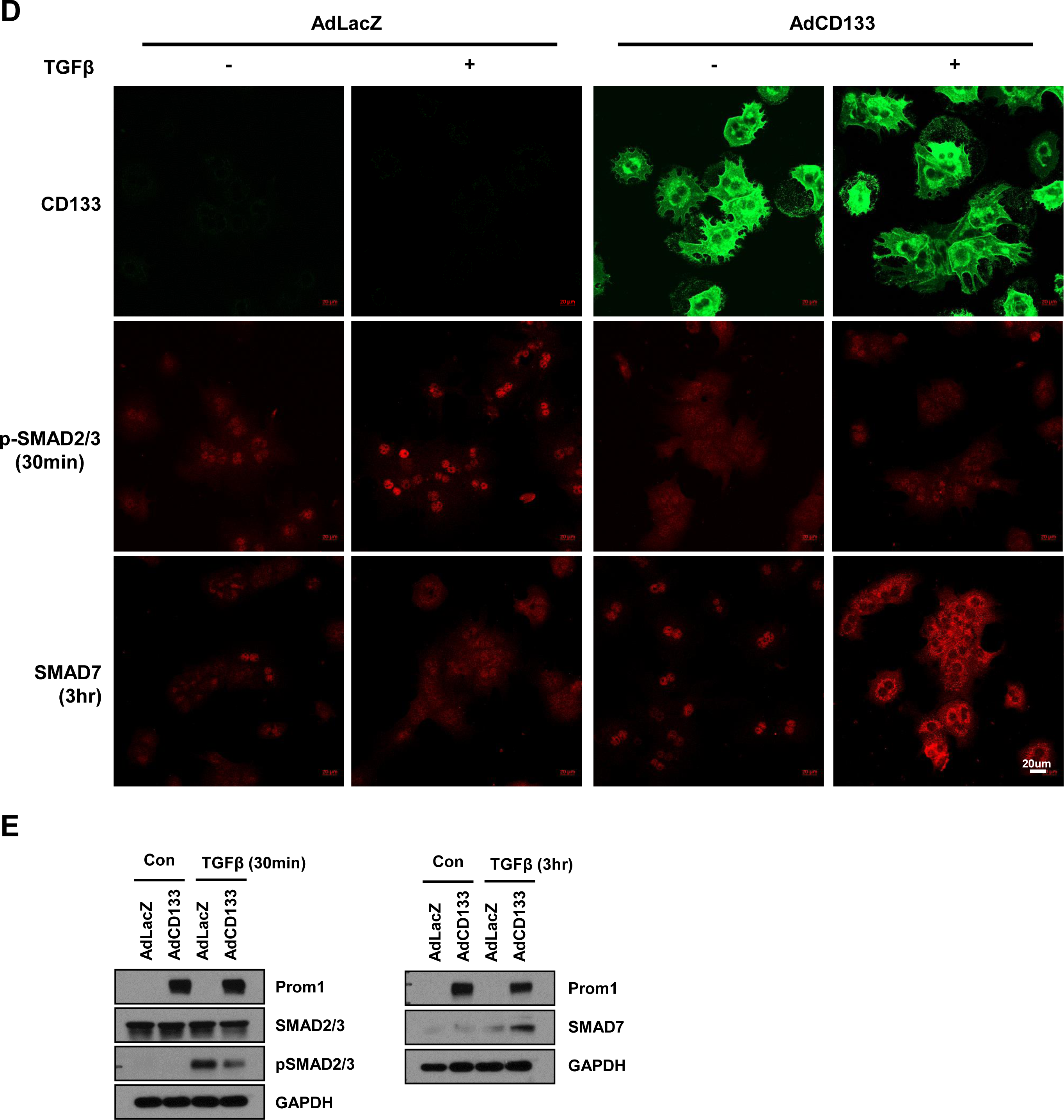
Prom1 deficiency enhances TGFβ signaling in hepatocytes. Primary hepatocytes were isolated from eight-week-old male *Prom1*^+/+^ and *Prom1*^-/-^ mice and grown for 4 h, serum-starved for 16 h and treated with 5 ng/ml TGF-β for the indicated times. The expression levels of PROM1, phosphorylated SMAD2/3 (p-SMAD2/3), total SMAD2/3 (tSMAD2/3), SMAD7 and Vimentin were determined by immunofluorescence and/or immunoblot analysis (A and B). Statistical analyses of protein expression after normalization to GAPDH were performed (B, right panel). After TGFβ treatment for 24 h, the mRNA levels of *Snail1*, *Vimentin*, *Zeb,1* and *Zeb2* were determined by qRT-PCR (C). Primary hepatocytes were isolated from *Prom1*^-/-^ mice, grown for 4 h, infected with adenovirus carrying LacZ or PROM1 for 24 h, serum-starved for 16 h and treated with 5 ng/ml TGFβ for the indicated time. The expression levels of PROM1, phosphorylated SMAD2/3 (pSMAD2/3), total SMAD2/3 (tSMAD2/3), SMAD7 and Vimentin were determined by immunofluorescence and/or immunoblot analyses (D and E). \**p* < 0.05, \*\**p* < 0.01.

### PROM1 stabilizes the SMAD7 protein

To elucidate how PROM1 stabilizes SMAD7, we examined the molecular interaction between PROM1 and SMAD7. A reciprocal endogenous immunoprecipitation assay in the BDL-treated Prom1^+/+^ liver showed a molecular interaction between PROM1 and SMAD7 (Fig. 6A). The interaction of PROM1 with SMAD7 was further demonstrated by exogenous immunoprecipitations in HEK 293T cells overexpressing both PROM1 and SMAD7 (Fig. 6B). To identify the domains of each protein that are necessary for the interaction, various truncation mutants of both PROM1 and SMAD7 were prepared (Supplementary Fig. 4A and B). Co-immunoprecipitations of both PROM1 and SMAD7 mutants showed that the 1^st^ intracellular domain (IC1) of PROM1 interacted with the N-terminal (N) domain or Mad homology 2 (MH2) domain of SMAD7 because all PROM1 mutants with the IC1 domain interacted with all SMAD7 mutants (Fig. 6C and D). To determine whether Prom1 stabilizes SMAD7, we overexpressed different combinations of PROM1, SMAD7 and SMURF1 and SMURF2 in HEK 293T cells and monitored SMAD7 protein expression by immunoblotting. The SMAD7 protein levels were decreased by SMURF2 but restored by Prom1 in a dose-dependent manner (Fig. 6E and F). In addition, SMAD7 ubiquitination which was induced by SMURF2 was abolished by PROM1 (Fig. 6G). The half-life of SMAD7 was determined by treating cells with cycloheximide (CHX). The half-life of SMAD7 in the presence of SMURF2 was increased 2-fold by PROM1 (Fig. 6H). To understand how PROM1 increases SMAD7 stability, we examined whether PROM1 and SMURF2 compete for interaction with SMAD7. Co-immunoprecipitation experiments showed that the molecular interaction between SMAD7 and SMURF2 was disrupted by PROM1 in a dose-dependent manner (Fig. 6I). Together, these data indicate that PROM1 abolishes SMURF2-induced SMAD7 ubiquitination by interfering with the molecular interaction between SMAD7 and SMURF2.

**Figure 6.**
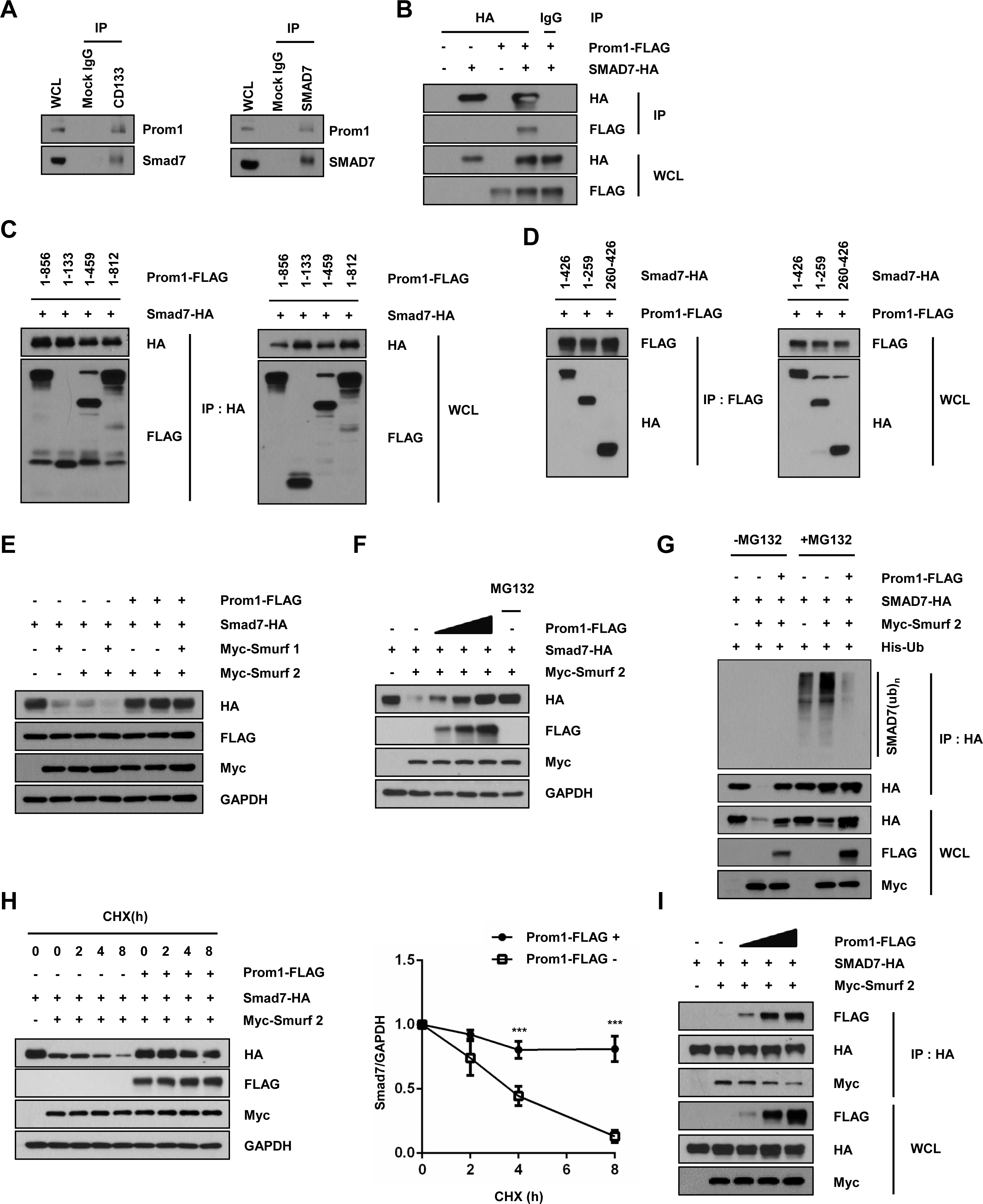
Prom1 prevents SMURF2-induced SMAD7 ubiquitination. (A) The molecular interaction between PROM1 and SMAD7 was assessed in *Prom1*^+/+^ hepatocytes by co-immunoprecipitation. (B) HEK293T cells were transfected with PROM1-FLAG and SMAD7-HA. The molecular interaction between PROM1-FLAG and SMAD7-HA was determined by co-immunoprecipitation (C and D) Different truncated mutants of PROM1-FLAG and SMAD7-HA were prepared and the indicated combinations were then transiently expressed in HEK 293T cells. Domain analysis was perfomed by co-immunoprecipitation with anti-HA (C) and anti-FLAG antibodies (D). (E) After transient expression of the indicated combinations of SMAD7-HA, Myc-SMURF-1, Myc-SMURF-2, and PROM1-FLAG in HEK 293T cells, the expression levels of SMAD7-HA, PROM1-FLAG, Myc-SMURF-1 and 2, and GAPDH were determined by immunoblotting. (F) Different amounts of PROM1-FLAG (0, 2, and 4 μg) were transiently expressed along with SMAD7-HA and Myc-SMURF2 in HEK 293T cells. The expression levels of SMAD7-HA, PROM1-FLAG and Myc-SMURF2 were determined by immunoblotting. MG132 treatment was used for a positive control for SMAD7-HA. (G) SMURF2-induced SMAD7 ubiquitination was decreased by PROM1 in HEK 293T cells. The indicated combinations of PROM1-FLAG, SMAD7-HA, Myc-SMURF2, and His-ubiquitin (His-Ub) were co-transfected into HEK 293T. After MG132 treatment, SMAD7 ubiquitination was determined by HA immunoprecipitation. (H) The degradation rate of SMAD7 protein expression with or without PROM1 was determined by immunoblotting. HEK 293T cells were co-transfected with the indicated combinations of PROM1-FLAG, SMAD7-HA, and Myc-SMURF2 and treated with 2 ug/ml cycloheximide (CHX) for the indicated times (left panel). The relative protein expression of SMAD7, shown in the left panel, was statistically calculated from three independent experiments (right panel). (I) PROM1 competes with SMURF2 for binding to SMAD7. HEK 293T cells were co-transfected with SMAD7-HA, Myc-SMURF2 and different amount of PROM1-FLAG (0, 2, and 4 μg). The molecular interaction of SMAD7-HA with Myc-SMURF2 and PROM1-FLAG was assessed by co-immunoprecipitation. WCL, whole-cell lysates; IP, immunoprecipitation. \*\*\**p* < 0.001.

### Liver-specific Prom1 deficiency aggravated BDL-induced liver fibrosis with reduced levels of SMAD7

To elucidate the specificity of hepatocytic PROM1 in liver fibrosis, we generated liver-specific *Prom1* deficient mice (*f/f : Alb-Cre*) by crossing *Prom1*^f/f^ mice (*f/f*), containing two loxP sequence flanking exon 2 of *Prom1* with Alb-Cre transgenic mice (*Alb-Cre*) (Supplementary Fig. 5A). PCR-based genotype of each mice were determined from the tail-tip of wild type, *Prom1*^f/f^ (*f/f*) and *Prom1*^f/f : Alb-Cre^ (*f/f : Alb-Cre*) mice (Supplementary Fig. 5B and C). Liver-specific Prom1 deficiency was confirmed using primary hepatocytes, liver and kidney of *Prom1*^f/f:Alb-Cre^ mice and its littermate *Prom1*^f/f^ mice by immunoblotting (Supplementary Fig. 5D).

Next we analyzed BDL-induced liver fibrosis in *Prom1*^f/f^ and *Prom1*^f/f : Alb-Cre^ mice. H&E, Sirius red and Masson’s trichrome staining assays revealed that *Prom1*^f/f : Alb-Cre^ mice aggravated BDL-induced liver fibrosis compared to Prom1^f/f^ mice (Supplementary Fig. 5E). BDL-induced expression of αSMA and CK19 was more increased in *Prom1*^f/f : Alb-Cre^ mice than in control *Prom1*^f/f^ mice as determined by qRT-PCR, immunofluorescence and immunoblotting, indicating that liver-specific *Prom1* deficiency aggravated BDL-induced liver fibrosis (Fig. 7A-C). Immunofluorescence and immunoblotting shows that BDL-induced phosphorylation of SMAD2/3 was also more increased with reduced levels of Smad7 caused by liver-specific *Prom1* deficiency (Fig. 7A and B and Supplementary Fig. 5E and 5G). However, increase of *Smad7* mRNA level after BDL did not differ, regardless of liver-specific knockout of *Prom1* (Supplementary Fig. 5F). With all these data, we conclude that hepatocytic PROM1 plays a central role in the regulation of TGFβ signaling and liver fibrosis via SMAD7 stabilization.

**Figure 7.**
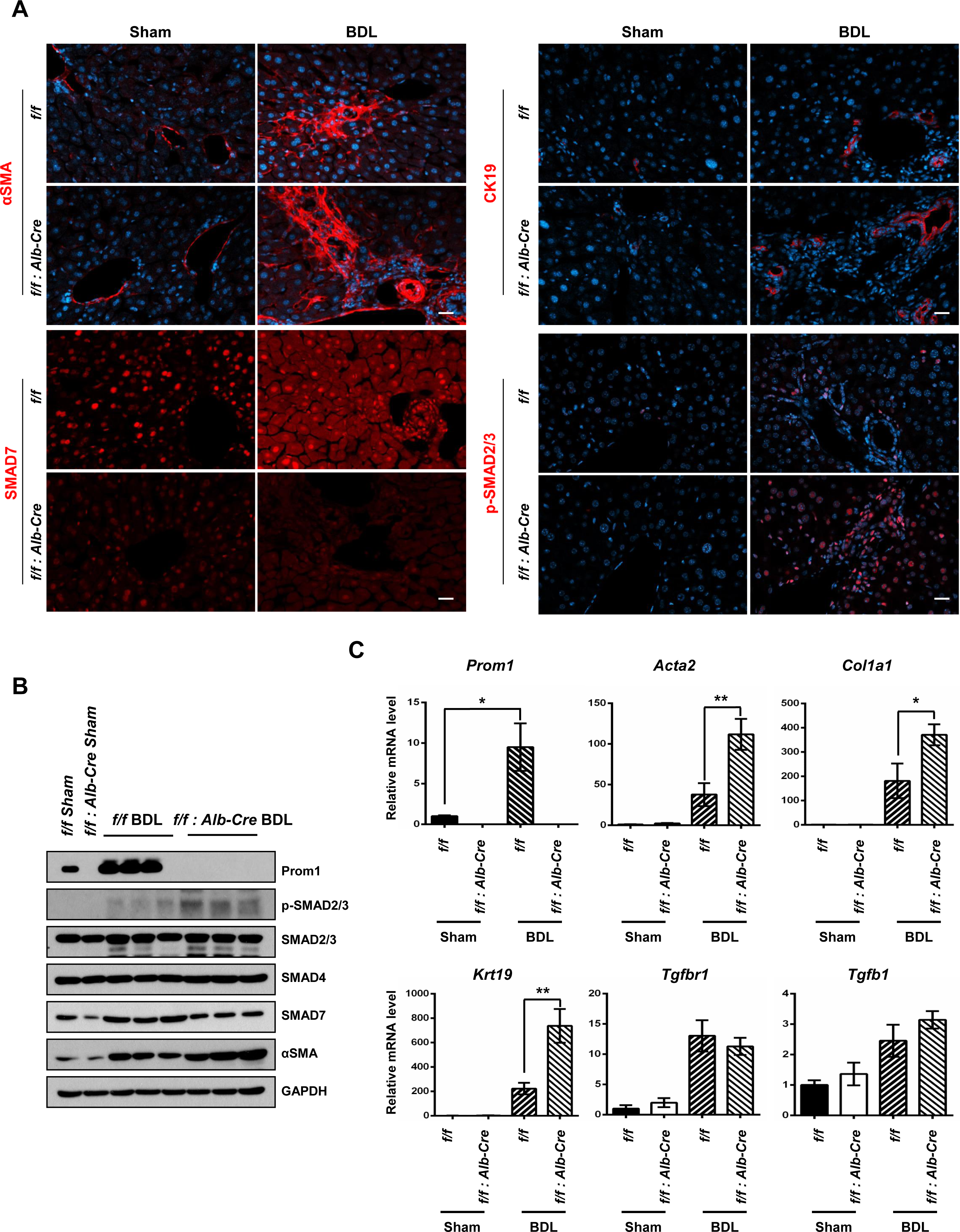
Liver specific PROM1 deficiency aggravates BDL-induced liver fibrosis. For BDL-induced liver fibrosis model, eight-week old male *Prom1*^f/f^ (f/f) and *Prom1*^f/f : Alb-Cre^ (f/f : Alb-Cre) mice were subjected to sham (n=3) or BDL (n=4-5) treatment for one week. Each liver specimen was analyzed by αSMA, CK19, SMAD7 and p-SMAD2/3 immunofluorescence (A). The expression levels of PROM1, phosphorylated SMAD2/3 (p-SMAD2/3), total SMAD2/3 (tSMAD2/3), SMAD4, SMAD7, αSMA and GAPDH were determined by immunoblot analysis (B). The mRNA levels of *Prom1, Acta2, Col1a1, Krt19, Tgfbr1,* and *Tgfb1* were determined by qRT-PCR and normalized to those of 18S rRNA (C). \**p* < 0.05, \*\**p* < 0.01

## Discussion

PROM1 was highly expressed in the fibrotic livers of human patients and mice (Fig. 1 and 2). To determine whether upregulated PROM1 expression is an important regulator or a simple result of liver fibrosis, we analyzed liver fibrosis in BDL- or CCl_4_-treated *Prom1*^+/+^ and *Prom1*^-/-^ mice. Because Prom1 deficiency aggravated BDL- or CCl_4_-induced liver fibrosis, Prom1 was determined as a negative regulator of liver fibrosis induced by TGFβ signaling. In our model based on the results, Prom1 negatively regulates TGFβ signaling in hepatocytes. PROM1 interacts with SMAD7 and increases SMAD7 protein expression by interfering with the molecular association of SMAD7 with SMURF2 and then preventing SMAD7 ubiquitination and degradation. In turn, SMAD7, being stabilized by PROM1, blocks the TGFβ-induced phosphorylation of SMAD2/3. Thus, PROM1 deficiency aggravates liver fibrosis by enhancing TGFβ signaling by stabilizing SMAD7 in hepatocytes (Figure 8).

**Figure 8.**
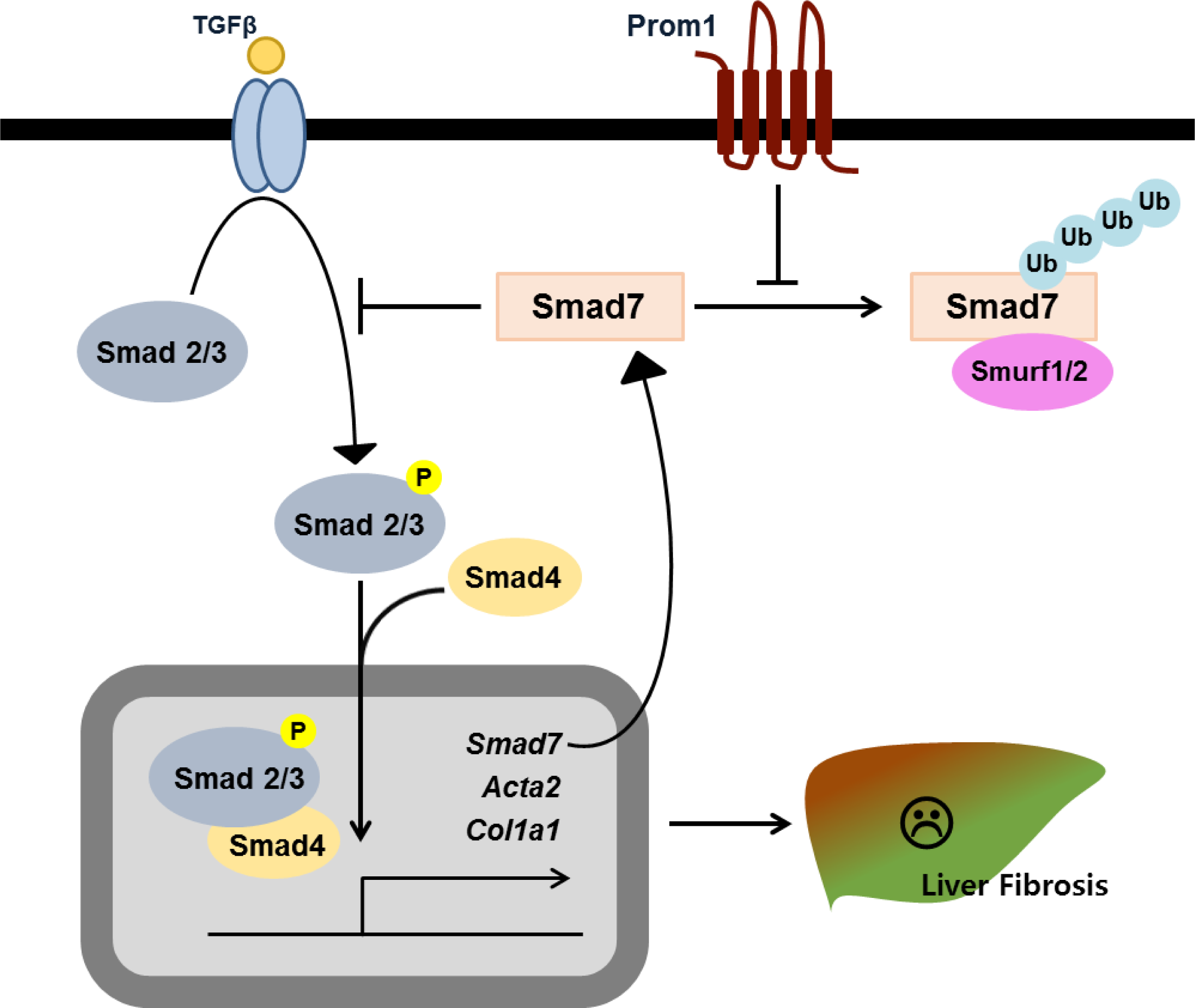
Graphical scheme explaining how PROM1 stabilizes the SMAD7 protein and protects the liver against liver fibrosis.

The numbers of Prom1- and CK19-expressing cells (HPCs and cholangiocytes) were significantly increased by BDL, which initiates a ductular reaction, but were marginally by CCl_4_ treatment, which damages mainly hepatocytes ^23, 24^. Interestingly, BDL-treated *Prom1*^-/-^ mice exhibited more CK19-expressing cells than to *Prom1*^+/+^ mice, suggesting that PROM1 is not necessary for the proliferation of CK19-expressing cells during liver fibrosis. Recent reports also show that PROM1 does not affect cholangiocyte proliferation in cholangiocyte-derived liver organoid ^25, 26^. Thus, PROM1 upregulation in the fibrotic liver of human patients and mice is the simple consequence of cholangiocyte proliferation.

Determining the tissue distribution and subcellular localization of PROM1 is difficult because of the limited immunoreactivity of glycosylated PROM1. To overcome this difficulty, we modified Shmelkov’s method to visualize PROM1 via immunofluorescence analysis ^27^. Our data shows that PROM 1 was localized in the plasma membrane of hepatocytes as well as in HPCs and cholangiocytes (Fig. 1C and 2B and 2D). Because the signal was not detected in the *Prom1*^-/-^ liver, the observed immunofluorescence signal was specific for PROM1. This observation allows us to focus on the role of hepatocytic PROM1 in liver fibrosis. Prom1^-/-^ hepatocytes showed increased TGFβ signaling than Prom1^+/+^ hepatocytes. For example, PROM1 deficiency increased TGFβ-induced phosphorylation of SMAD2/3, and the transcription of *Snail1*, *Vimentin*, *Zeb1,* and *Zeb2* while reducing SMAD7 protein expression. Thus, we conclude that Prom1 deficiency in aggravates BDL- and CCl_4_-induced liver fibrosis.

Recently, Zagory *et al.* demonstrated that Prom1 promotes RRV-induced biliary fibrosis ^22^. They showed that *Prom1* disruption prevented RRV-induced biliary fibrosis by assessing fibrosis with Sirius Red staining, immunofluorescence and qRT-PCR of CK19 and INTEGRIN-β6. However, serum assays of bilirubin did not show that Prom1 disruption prevented RRV-induced biliary fibrosis. Unlike theirs, our results indicated that Prom1 disruption aggravated BDL- or CCl_4_-induced liver fibrosis by increasing SMAD7 degradation.

PROM1 is an important surface marker of normal and cancer stem cells and has several binding partners such as Actin, Radixin, PI3K, and HDAC6^28–31^. In addition to these binding partners, we demonstrated that PROM1 plays a central role in TGFβ signaling and liver fibrosis by stabilizing SMAD7 using both global and liver-specific Prom1 knockout mice. We believe that our study will shed light on a novel physiological function of PROM1 in liver fibrosis. In addition to the liver, fibrosis progresses in other organs such as the heart, lung, kidney, and skin where PROM1 is highly expressed ^17, 32, 33^. Because the role of SMAD7 during fibrosis progression has been extensively studied in these organs ^34–37^, a PROM1-SMAD7 axis will provide a better insight into understanding fibrosis in other organs.

---

Supplementary Figure 1. Prom1 deficiency aggravates BDL- and CCl_4_-induced liver fibrosis. For BDL-induced liver fibrosis model, eight-week old male *Prom1*^+/+^ and *Prom1*^-/-^ mice were subjected to sham (n=3) or BDL (n=5-6) treatment for one week. For the CCl_4_-induced liver fibrosis model, 8-week old male *Prom1*^+/+^ and *Prom1*^-/-^ mice were administrated with intraperitoneal injections of vehicle (n=6) or CCl_4_ (n=12-13) twice a week for 6 weeks. Each liver specimen was analyzed by Prom1 immunofluorescence and H&E staining (A). The mRNA levels of *Ctgf, Pai1, Timp1, Il1b, Il6, Tnfa,* and *Ccl2* were determined by qRT-PCR and normalized to those of 18S rRNA (B). \**p* < 0.05, \*\*\**p* < 0.001.

Supplementary Figure 2. PROM1 deficiency does not alter *Smad7* mRNA expression in mice with liver fibrosis. For the BDL-induced liver fibrosis model, eight-week old male *Prom1*^+/+^ and *Prom1*^-/-^ mice were subjected to sham or BDL treatment for one week. For CCl_4_-induced liver fibrosis model, 8-week old male *Prom1*^+/+^ and *Prom1*^-/-^ mice were administrated with intraperitoneal injections of vehicle or CCl4 twice a week for 6 weeks. The mRNA levels of *Smad7* were determined by qRT-PCR and normalized to those of 18S rRNA (A). The protein expression levels of αSMA were normalized to that of GAPDH and statistically assessed. N=3 for each group (B). \**p* < 0.05.

Supplementary Figure 3. Prom1 deficiency does not alter *Smad7* mRNA expression in primary mouse hepatocytes. Primary hepatocytes were isolated from 8-week-old male *Prom1*^+/+^ and *Prom1*^-/-^ mice and grown for 4 h, serum-starved for 16 h and treated with 5 ng/ml TGF-β for the indicated times. The localization of phosphorylated SMAD2/3 was determined by immunofluorescence, as shown in Fig. 5A. The percentage of cells with nuclear p-SMAD2/3 expression, shown in Fig. 5A images (A), was statistically assessed. After TGFβ treatment for 24 h, the mRNA levels of *Prom1* and *Smad7* were determined by qRT-PCR. \**p* < 0.05.

Supplementary Figure 4. (A and B) Structure of *PROM1* (A) and *SMAD7* (B) truncation mutants used in domain analysis. EX, extracellular domain; TM, transmembrane domain; IC, intracellular domain; N domain, N-terminal domain; MH2 domain, Mad homology 2 domain.

Supplementary Figure 5. Liver specific PROM1 deficiency aggravates BDL-induced liver fibrosis. Schematic for generating liver-specific *Prom1* deficient mice (A). Schematic representation of the genomic structures near the second exon of *Prom1* (WT*, f/f*) and the promoter region of *Alb* (*Alb-Cre*). Mice were created by crossing *Prom1^f/f^*mice (*f/f*), containing two loxP sequences flanking exon 2 of *Prom1*, with Alb-Cre transgenic mice (*Alb-Cre*). Areas targeted by primers used for genotyping are marked in B. Representative PCR products from tail-tip genomic DNA of WT, Prom1^loxp/loxp^ (f/f) and Prom1^loxp/loxp^ /Alb-Cre (f/f : Alb-Cre) mice (B and C). The expression levels of PROM1 in primary hepatocytes, liver and kidney of *f/f* and *f/f : Alb-Cre* were determined by immunoblot analysis (D). For BDL-induced liver fibrosis model, eight-week old male Prom1^loxp/loxp^ (*f/f*) and Prom1^loxp/loxp^ /Alb-Cre (*f/f : Alb-Cre*) mice were subjected to sham (n=3) or BDL (n=4-5) treatment for one week. Each liver specimen was analyzed by PROM1 immunofluorescence, Masson’s Trichrome staining, Sirius Red staining and H&E staining (E). The mRNA levels of *Smad7* were determined by qRT-PCR and normalized to those of 18S rRNA (F). The expression levels of pSMAD2/3 and SMAD7 were normalized to that of GAPDH and statistically assessed (G). N=1 for sham group and N=3 for BDL group. \**p* < 0.05, \*\**p* < 0.01.

Supplementary Table 1. List of antibodies used in this study.

Supplementary Table 2. List PCR primers used in this study.

## Notes

**Grant support** This work was supported by grants awarded to Y.-G. Ko, H. Lee, and D.-M. Yu from the National Research Foundation (R1A5A1009024, R1912691 and R1821961).

**Conflicts of interest** The authors disclose no conflicts

#### Summary of Updates

Title is revised (Hepatocytic Prominin-1 protects liver fibrosis by stabilizing the SMAD7 protein -> Hepatocytic Prominin-1 protects against liver fibrosis by stabilizing the SMAD7 protein) Additional author

