## supplementary figure 1-5, supplementary table 1-2 for "Hepatocytic Prominin-1 protects against liver fibrosis by stabilizing the SMAD7 protein"

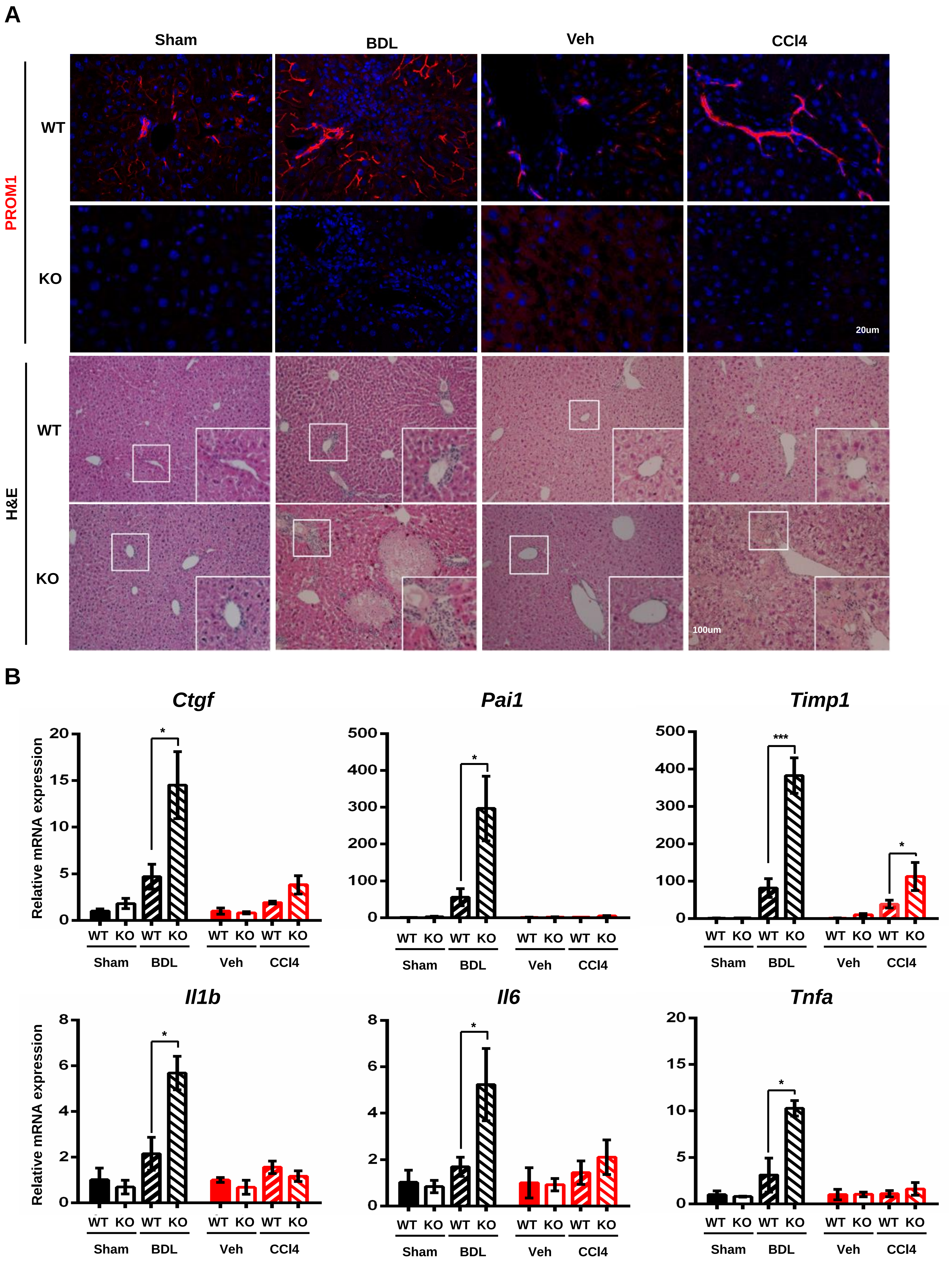


**Supplementary Figure 1**

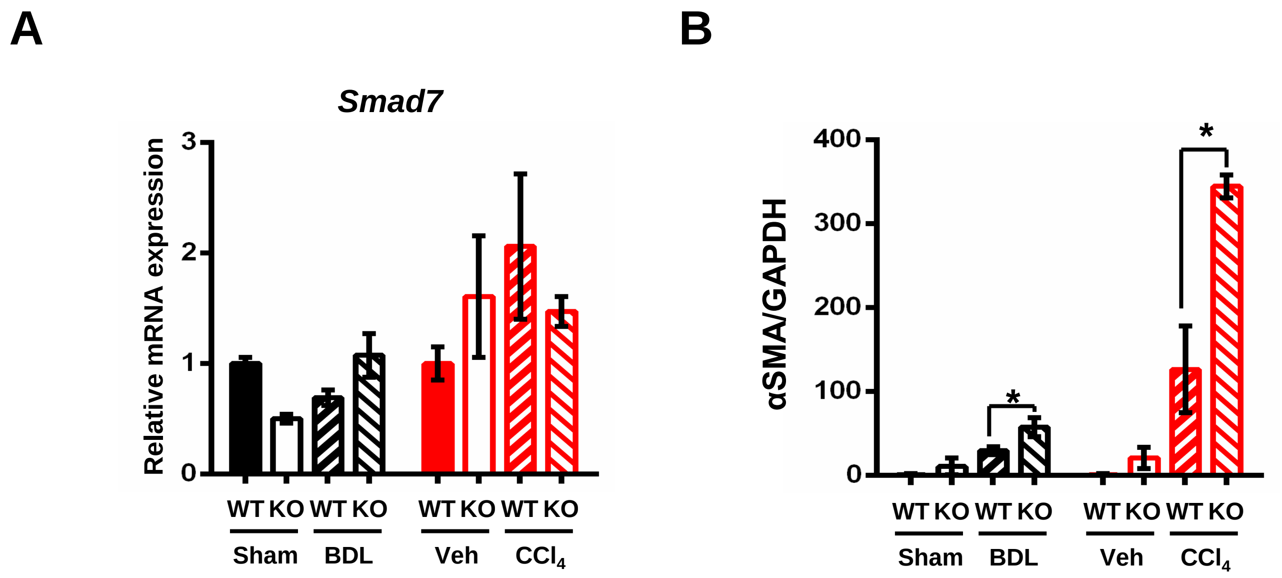


**Supplementary Figure 2**

A

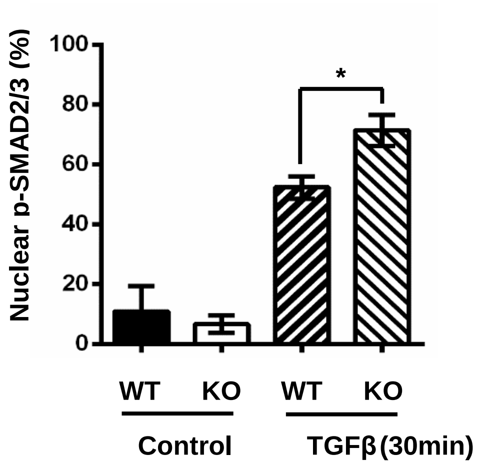


B

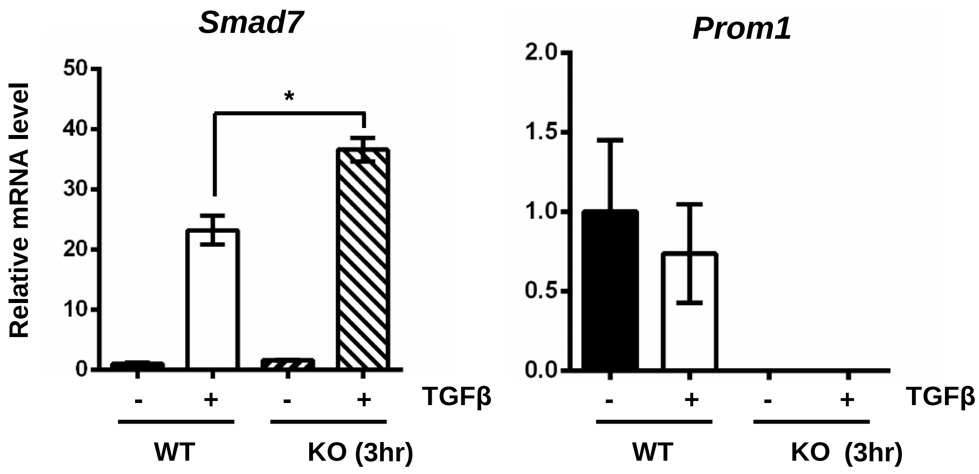


Supplementary Figure 3

**A**

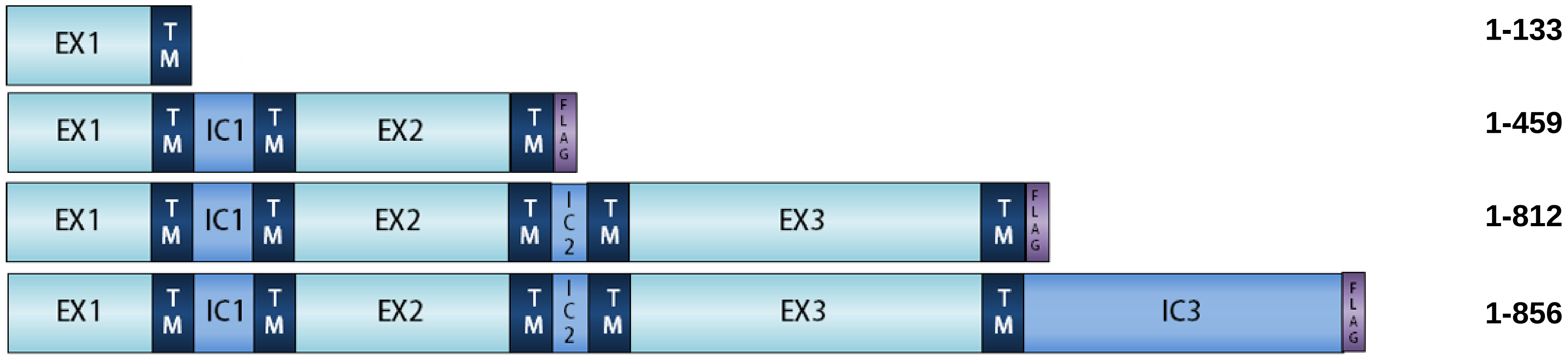


**B**

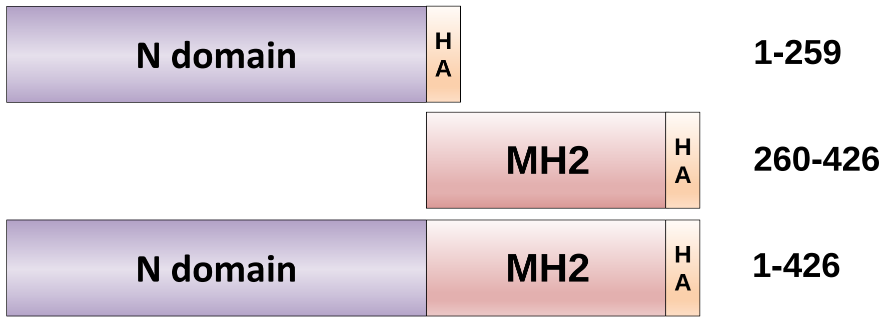


**Supplementary Figure 4**

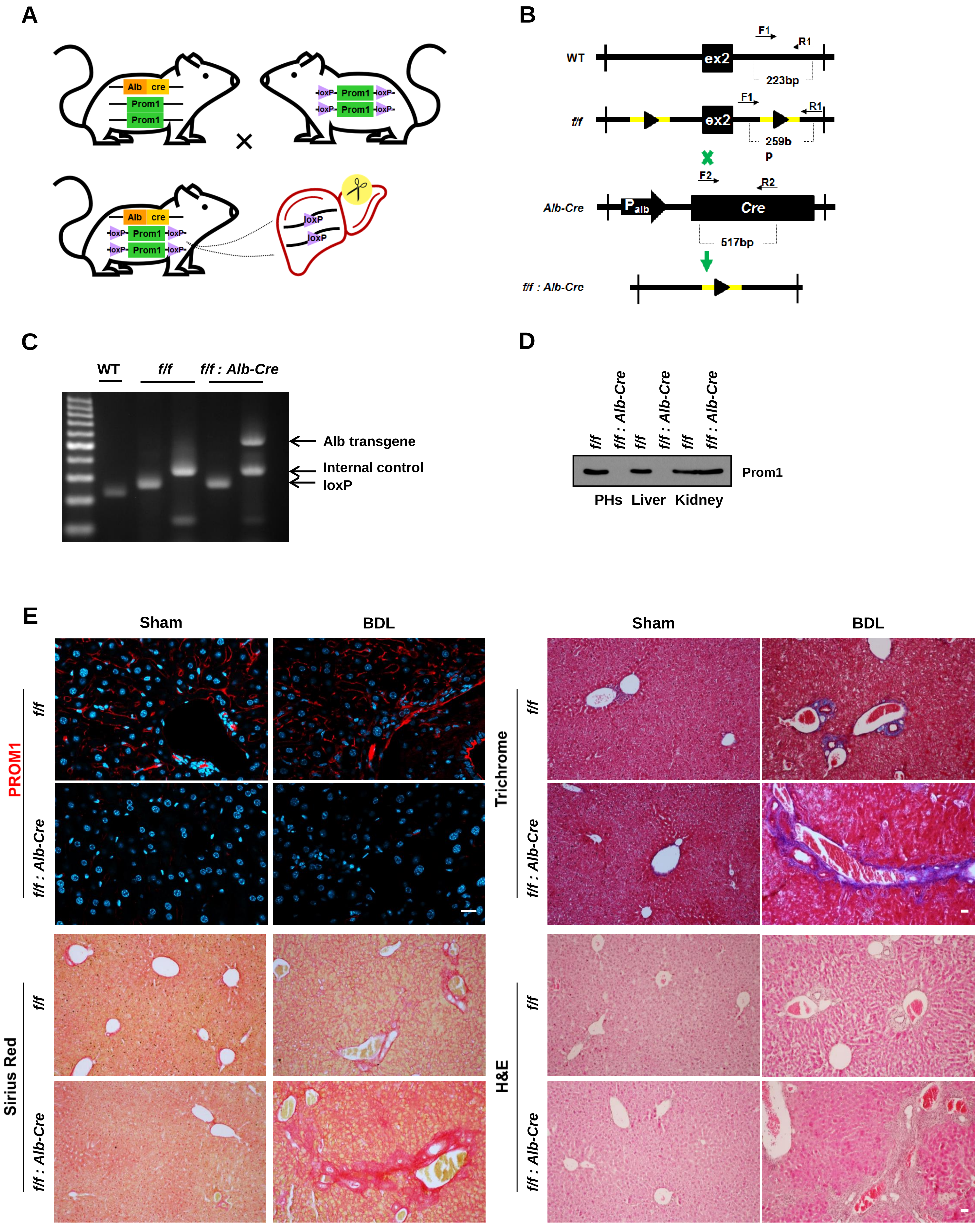


**Supplementary Figure 5**

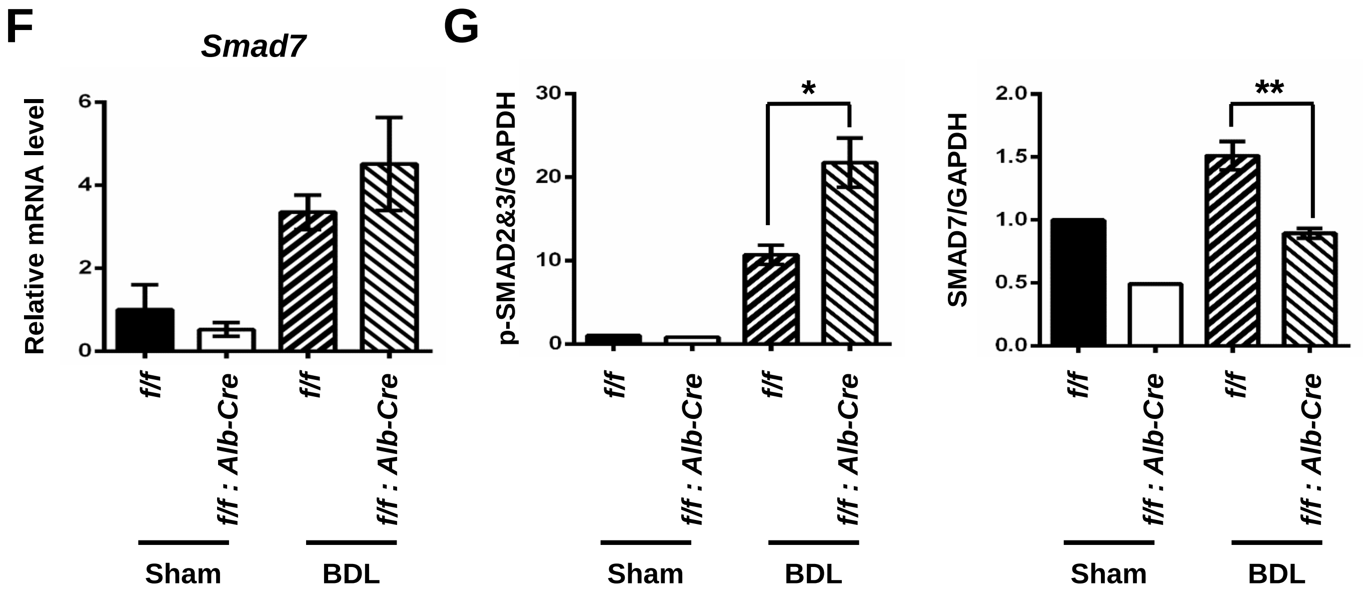


Supplementary Figure 5

| Name | Company | Host | Experiments |
|---|---|---|---|
| Prom1 | eBioscience | Rat polyclonal | IB(1:2000), IHC(1:100) |
| Prom1 | MACS | Mouse Monoclonal | IB(1:200), IFA(1:25) |
| Prom1 | Merck Millipore | Mouse monoclonal | IB(1:1000), IFA(1:100) |
| Prom1 | R&D systems | Mouse monoclonal | IB(1:1000), IFA(1:100) |
| α-SMA | abcam | Rabbit polyclonal | IB(1:1000),IHC(1:200) |
| CK19 | abcam | Mouse Monoclonal | IB(1:1000),IHC(1:200) |
| SMAD2/3 | Cell Signaling Technology | Rabbit polyclonal | IB(1:1000) |
| pSMAD2/3 | Cell Signaling Technology | Rabbit polyclonal | IB(1:500), IFA(1:100) |
| SMAD4 | Cell Signaling Technology | Rabbit polyclonal | IB(1:1000) |
| SMAD7 | R&D systems | Mouse monoclonal | IB(1:1000), IFA(1:100) |
| Vimentin | abcam | Rabbit monoclonal | IB(1:1000), IFA(1:100) |
| Flag | Sigma-Aldrich | Rabbit monoclonal | IB(1:2000) |
| HA | Merck Millipore | Mouse monoclonal | IB(1:1000) |
| Myc | Santa Cruz Biotechnology | Mouse monoclonal | IB(1:1000) |
| GAPDH | Santa Cruz Biotechnology | mouse monoclonal | IB(1:2000) |

**Supplementary Table 1**

| | Name | Forward | Reverse |
|---|---|---|---|
| qRT-PCR | Prom1 | CTCATGGCTGGGGTTGGATT | TGAGCAGATAGGGAGTGTCCA |
| | Acta2 | AGCTACGAACTGCCTGACGG | CGTGGATGCCCGCTGAC |
| | Col1a1 | AGCACGTCTGGTTTGGAGAG | GACATTAGGCGCAGGAAGGT |
| | Tgfb1 | GTCACTGGAGTTGTACGGCA | GGGGCTGATCCCGTTGATTT |
| | Tgfbr1 | GCATTGGCAAAGGTCGGTTT | TGCCTCTCGGAACCATGAAC |
| | Krt19 | GTGCTGGATGAGCTGACTCTG | GATCTTGGCTAGGTCGACACC |
| | Ctgf | AGGGCCTCTTCTGCGATTTC | CTTTGGAAGGACTCACCGCT |
| | Pai1 | CACAGGCACTGCAAAAGGTC | TGTGCCGAACCACAAAGAGA |
| | Timp1 | GGCATCTGGCATCCTCTTGT | TGGTCTCGTTGATTTCTGGGG |
| | Il1b | GCCCATCCTCTGTGACTCAT | AGGCCACAGGTATTTTGTCG |
| | Il6 | TGATGGATGCTACCAAACTGGA | ACTCTGGCTTTGTCTTTCTTGT |
| | Tnfa | TGGGACAGTGACCTGGACTGT | TTCGGAAAGCCCATTTGAGT |
| | Smad7 | GGGGGCTTTCAGATTCCCAA | GACACAGTAGAGCCTCCCCA |
| | Snail1 | GTCTGCACGACCTGTGGAAA | GGTCAGCAAAAGCACGGTTG |
| | Vimentin | TTCTCTGGCACGTCTTGACC | GCTTGGAAACGTCCACATCG |
| | Zeb1 | TGGCAAGACAACGTGAAAGA | AACTGGGAAAATGCATCTGG |
| | Zeb2 | TAGCCGGTCCAGAAGAAATG | GGCCATCTCTTTCCTCCAGT |
| Genotype PCR | Cre transgene | CCAGGCTAAGTGCCTTCTCTACA | AATGCTTCTGTCCGTTTGCCGGT |
| | Internal control | CTAGGCCAGAGAATTGAAAGATCT | GTAGGTGGAAATTCTAGCATCATCC |
| | loxP | ATGGGGAGCATTGATTTGCC | TCTTTCTGCCTGGTTGCTTT |

Supplementary Table 2
